# CSPα reduces aggregates and rescues striatal dopamine release in αsynuclein transgenic mice

**DOI:** 10.1101/2020.07.31.229153

**Authors:** L Caló, E Hidari, M Wegrzynowicz, JW Dalley, BL Schneider, O Anichtchik, E Carlson, D Klenerman, MG Spillantini

## Abstract

αSynuclein aggregation at the synapse is an early event in Parkinson’s disease and is associated with impaired striatal synaptic function and dopaminergic neuronal death. The cysteine string protein (CSPα) and αsynuclein have partially overlapping roles in maintaining synaptic function and mutations in each cause neurodegenerative diseases. CSPα is a member of the DNAJ/HSP40 family of co-chaperones and like αsynuclein, chaperones the SNARE complex assembly and neurotransmitter release. αSynuclein can rescue neurodegeneration in CSPαKO mice. However, whether αsynuclein aggregation alters CSPα expression and function is unknown. Here we show that αsynuclein aggregation at the synapse induces a decrease in synaptic CSPα and a reduction in the complexes that CSPα forms with HSC70 and STGa. We further show that viral delivery of CSPα rescues *in vitro* the impaired vesicle recycling in PC12 cells with αsynuclein aggregates and *in vivo* reduces synaptic αsynuclein aggregates restoring normal dopamine release in 1-120hαsyn mice. These novel findings reveal a mechanism by which αsynuclein aggregation alters CSPα at the synapse, and show that CSPα rescues αsynuclein aggregation-related phenotype in 1-120hαsyn mice similar to the effect of αsynuclein in CSPαKO mice. These results implicate CSPα as a potential therapeutic target for the treatment of early-stage PD.

## Introduction

Alpha-synuclein (αsyn) is a synaptic protein involved in vesicle clustering, assembly of the SNARE complex and neurotransmitter release. Point mutations and duplication/triplication of the αsynuclein gene cause Parkinson’s disease (PD) (Lunati *et al*., 2018) and αsyn aggregates form the Lewy bodies characteristic of PD (Spillantini *et al*., 1998, 1997). C-terminal truncation of αsyn, found in Lewy bodies, promotes its aggregation (Baba *et al*., 1998; Crowther *et* al., 1998). We previously described 1-120hαsyn transgenic mice expressing C-terminally truncated αsyn under the control of the tyrosine hydroxylase-promoter in the absence of the endogenous protein (Tofaris *et al*., 2006), where αsyn aggregation in the striatal terminals is associated with re-distribution of SNARE proteins and impairment in dopamine (DA) release, features present in PD patients (Garcia-Reitbock *et al*., 2010). Growing evidence points to presynaptic terminals as the initial site of neurodegeneration in PD (Nakata *et al*., 2012; Janezic *et al*., 2013; Garcia-Reitbock *et al*., 2010; Wegrzynowicz *et al*., 2019), as shown in our transgenic MI2 mice, where synaptic dysfunction with αsyn accumulation preceded DA cell death, both rescued by an oligomer modifier (Wegrzynowicz *et al*., 2019).

The cysteine string protein α (CSPα/DNAJC5) is a vesicle-associated protein that regulates neurotransmitter release, exocytosis/endocytosis coupling and SNARE complex assembly through a pathway parallel to that of αsyn (Tobaben *et al*., 2001; Chandra *et al*., 2005; Zhang *et al*., 2012), DNAJC proteins have been linked to parkinsonism (Roosen *et al*. 2019). CSPα function is mediated by the DNAJ domain which activates the ATPase activity of the Heat shock cognate 70kDa protein HSC70 (Braun *et al*., 1996; Chamberlain *et al*., 1997); its mechanism of action is functionally associated with αsyn in that overexpression of αsyn abolishes lethal neurodegeneration in CSPα-KO mice and ablation of all three (α,β,γ)-syn genes results in SNARE complex assembly deficit with an increase in CSPα (Burre *et a*l., 2010; Goremberg *et al*., 2017). However, whether CSPα levels and activity change with αsyn aggregation and SNARE protein redistribution or if CSPα can rescue the synaptic pathology associated with αsyn aggregation is not known.

In this study we show that CSPα expression and function are impaired in the presynaptic terminals of 1-120hαsyn mice concomitantly with the presence of αsyn aggregates and a reduction in evoked DA release. We further show that expression of CSPα rescues both αsyn -aggregation dependent deficit in vesicle cycle *in vitro* and impaired DA release *in vivo*. This effect is associated *in vivo* with reduction in the number of striatal synaptic αsyn aggregates as shown by super resolution analysis of tissue sections.

## Materials and Methods

### Mice

Transgenic 1-120hαsyn and control mice without endogenous αsyn (C57BL/6/OlaHsd) were used in this study (Tofaris *et al*., 2006; Garcia-Reitbock *et al*., 2010). Regulated animal procedure were carried out under the Animals (Scientific Procedures) Act 1986 Amendment Regulations 2012 following ethical review by the University of Cambridge Animal Welfare and Ethical Review Body (AWERB), under project license no. 7008383.

### Immunostaining

Brains from paraformaldehyde-perfused 12 month-old mice were sectioned and 30 µm free-floating sections were incubated overnight at 4°C with primary antibodies (anti-αsyn, BD Transduction, 1:700), anti-DNAJC5 (Millipore, 1:500) as previously done (Garcia-Reitbock *et al*., 2010). Staining was visualised using the ABC Elite Kit (Vector Laboratories) and 3,3′-diaminobenzidine or Alexa-labelled secondary antibodies and imaged using a Leitz DMRB microscope or a Leica TCS SPE confocal microscope.

### Co-immunoprecipitation and Immunoblotting

Total proteins were extracted from mouse striata in PBS containing 0.1% Tween 20 and Protease Inhibitor cocktail (Roche) with or without 1 mM non-hydrolysable ADP (Sigma). Immunoprecipitation followed previous protocols (Garcia-Reitbock *et al*., 2010). Briefly, proteins (0.8-1 mg) were rotated overnight at 4°C with 5 µg of mouse anti-SGTa antibody (Abcam) or control mouse IgG and protein G Dynabeads (Invitrogen). Immunocomplexes were eluted by denaturation in NuPAGE LDS sample buffer (Invitrogen). Synaptosomal fractions were extracted using Syn-PER synaptic protein extraction reagent (Thermo Scientific). Proteins were resolved on a 4-12% gradient PAGE-SDS gel (Invitrogen), transferred onto nitrocellulose membranes (Bio-Rad), incubated with peroxidase-conjugated secondary antibodies (GE Healthcare) and visualised with chemiluminescent substrates (Thermo Fisher Scientific) as previously described (Wegrzynowicz *et al*., 2019). Antibodies were: mouse anti-DNAJC5 (Millipore1:500), anti-VAMP2 (Abcam 1:500), anti-ATPase HSC70 (Synaptic systems, 1:500), anti-SGTa (Abcam 1:500) and rabbit anti-β-actin (Abcam, 1:10000).

### AAV vector injections

Human full-length CSPα cDNA, a gift from Prof RD Burgoyne (Liverpool University), was subcloned into a pAAV vector under the PGK promoter and packaged in serotype 6 AAV particles as described (Löw *et al*., 2013). Vector suspension was diluted to 1×10^13^ viral genome containing particles (VG)/mL and an AAV6 empty vector (EV) used as control. For vector injections, animals were anesthetized with 2% isoflurane, placed in a stereotaxic frame (David Kopf Instruments) and injected bilaterally with 2 μl of the virus (0.2 μl/min flow rate) in the substantia nigra (SN) at the following coordinates: AP=±0.7, L=±1.7, DV=-3.6 below dural surface relative to the bregma according to Paxinos and Watson (Paxinos and Franklin, 2004).

### Retention of FM1-43

PC12 cells stably expressing 1-120hαSyn were plated onto coverslips in 12 well plates at 7×10^5^ cells/ml (Garcia-Reitbock *et al*., 2010). Cells were infected with 1μl 1×10^13^ VG/mL AAV6CSPα or control AAV6EV for 24 hours, then grown for 4 days in fresh medium. To stimulate vesicle endocytosis of the FM1-43 dye (Invitrogen), PC12 cells were depolarised with KCl (Hank’s balanced salts medium with Ca^2+^ and Mg^2+^, 90 mM KCl, 63 mM NaCl) then incubated with 15 µM FM1-43 for 90 s at room temperature and unbound dye removed by 10 min wash in PBS /1mM scavenger dye ADVASEP-7 (Sigma). Cells were re-incubated with depolarizing solution for 90 s at room temperature (Gaffield *et al*., 2006; Garcia-Reitbock *et al*., 2010). Only vesicles with impaired release retained the dye. Cells were then washed, fixed with 4% paraformaldehyde and stained (Syn1 antibody, BD Biosciences, 1:500) overnight at 4°C. Signal was detected using a Leica SPE 4 confocal microscope. FM1-43 positive puncta above the threshold fluorescence set by the AAV6EV transduced cells were counted using ImageJ analysis software. Between 700-1000 cells were counted for each experimental condition.

### *In vivo* microdialysis

*In vivo* microdialysis was performed as previously reported (Garcia-Reitbock *et al*., 2010; Wegrzynowicz *et al*., 2019). A microdialysis cannula (CMA Microdialysis) was placed in anesthetized mice in the right medial striatum (AP = ±0.7, L = ±1.7, H = -2.1 from the bone [31], DV = -2.1 from the skull surface). The following day, a CMA/7 microdialysis probe was inserted into the guide cannula and perfusion performed at a constant flow rate (2 µl/min) with artificial cerebrospinal fluid (ACSF: 140 mM NaCl, 7.4 mM glucose, 3 mM KCl, 0.5 mM MgCl_2_, 1.2 mM CaCl_2_, 1.2 mM Na_2_HPO_4_, 0.3 mM NaH_2_PO_4_, pH 7.4). Dialysates were collected every 20 min in tubes containing 5 µl of 0.2 M perchloric acid to prevent dopamine oxidation and assayed for dopamine, homovanillic acid and 3,4-dihydroxyphenylacetic acid. Two fractions (20-40 min) were collected to evaluate baseline release, ACSF was then replaced by ACSF containing 50 mM KCl and three more fractions collected (60-100 min). High KCl ACSF was then replaced by basal ACSF and two more fractions (120-140 min) were collected after which mice were killed, and brains used for immunohistochemistry or immunoblotting. Dopamine and homovanillic acid levels in the dialysate were measured by high-performance liquid chromatography (Garcia-Reitbock *et al*., 2010; Wegrzynowicz *et al*., 2019).

### *d*STORM

Thirty-micron free-floating striatal sections were stained for αsyn (Syn1, BD, 1:300) and Alexa Fluor Plus 647 secondary antibody (Invitrogen, 1:2000) in the presence of Tetra Speck microspheres (Invitrogen) to correct for drift during imaging.

Images were acquired with Photometrics EMCCD camera on a Nikon Ti-2E inverted microscope in near-TIRF mode. Image stacks consisted of 10,000 frames (50 ms/frame) on the field of view (FOV). A total of four-five FOVs per section and four mice per group were tested.

The image stacks were drift corrected and analysed using PeakFit in the open source ImageJ plugin GDSC SMLM, followed by a custom script. Briefly, this grouped the fluorescence signals into clusters (monomers or aggregates) based on their spatiotemporal distribution to determine their area. The aggregate size reported is the square root of the area (Whiten *et al*., 2018,*b*).

### Experimental design and statistical analysis

#### Immunoblotting

relative band intensity (RI) was calculated using Image J and βActin-normalised CSPα, HSC70, STGa and VAMP2 levels analysed with two-tailed Student’s t test.

#### Co-immunoprecipitation

RI of CSPα, HSC70, and SGTa alongside corresponding input proteins were normalised to βActin and analysed using one-way ANOVA with Bonferroni’s multiple comparisons test.

#### FM1-43 dye retention

data were evaluated using two-way ANOVA with Bonferroni’s multiple comparison test.

#### Microdialysis

DA release was normalized to the baseline fraction (0 min) and expressed as fold difference relative to the average DA release directly following K^+^ stimulation (60 min fraction) in the control group. DA release in 1-120hαSyn or control mice treated with CSPα or EV was analysed using two-way ANOVA with Bonferroni’s multiple comparison test.

#### *d*STORM

species were divided into monomer/aggregates based on their size with recombinant monomers having a median length < 36.5 nm (20.65±2×7.9 mean±2×SD, Wegrzynowicz *et al*., 2019). Data were analysed using the software https://github.com/Eric-Kobayashi/SR_toolkit.

Median size was analysed using a two-tailed Student’s t test; size distribution differences between CSPα and EV-treated groups (monomers and aggregates combined) was calculated from a cumulative histogram using a Kolmogorov-Smirnov test.

## Results

### CSPα levels are altered in the striatum of 1-120hαSyn mice

αSyn aggregates were present in the striatum of 1-120hαSyn 12 month-old mice, no αsyn staining was present in background control mice lacking the endogenous protein (Tofaris *et al*., 2006; Garcia-Reitbock *et al*., 2010). CSPα staining in the striatum of control mice revealed a punctate pattern that was less intense in 1-120hαSyn mice (Fig 1A). By immunoblotting no significant difference was present in CSPα amounts between controls and 1-120hαSyn mice in total tissue homogenates, however, a significant 68.3% reduction in CSPα amount was present in synaptosomal protein extracts in 1-120hαSyn mice (Fig. 1B) (Total homogenates RI: Control 1.29±0.26; 1-120hαSyn 1.07±0.27; Synaptic fraction: control 1.83±0.21; 1-120hαSyn 0.58±0.16). HSC70 and SGTa were not changed in the synaptic fraction (Supplementary Fig. 1). The level of the synaptic v-SNARE protein VAMP2 was also not changed as previously reported (Garcia-Reitbock *et al*., 2010; Supplementary Fig. 1). CSPα in synaptic terminals forms a trimeric complex with HSC70 and SGTa. To test whether this function was perturbed in the striatum of 1-120hαSyn mice, we used immunoprecipitation in the presence of ADP. We found a reduction in the amount of CSPα and HSC70 complexes in 1-120hαSyn mice compared to controls. No changes in SGTa, HSC70 and CSPα expression levels were found in the total homogenate input (Fig. 1C) (RI: HSC70: control 93031±3690, 1-120hαSyn 53078±1726; CSPα: control 69117±3181, 1-120hαSyn 18097±2762; SGTa: control 87661±5166, 1-120hαSyn 81385±4499). Taken together, these findings suggest that the presence of aggregated αsyn in 1-120hαSyn mice selectively reduces CSPα levels in striatal synaptic terminals hampering its chaperone activity.

**Figure 1.**
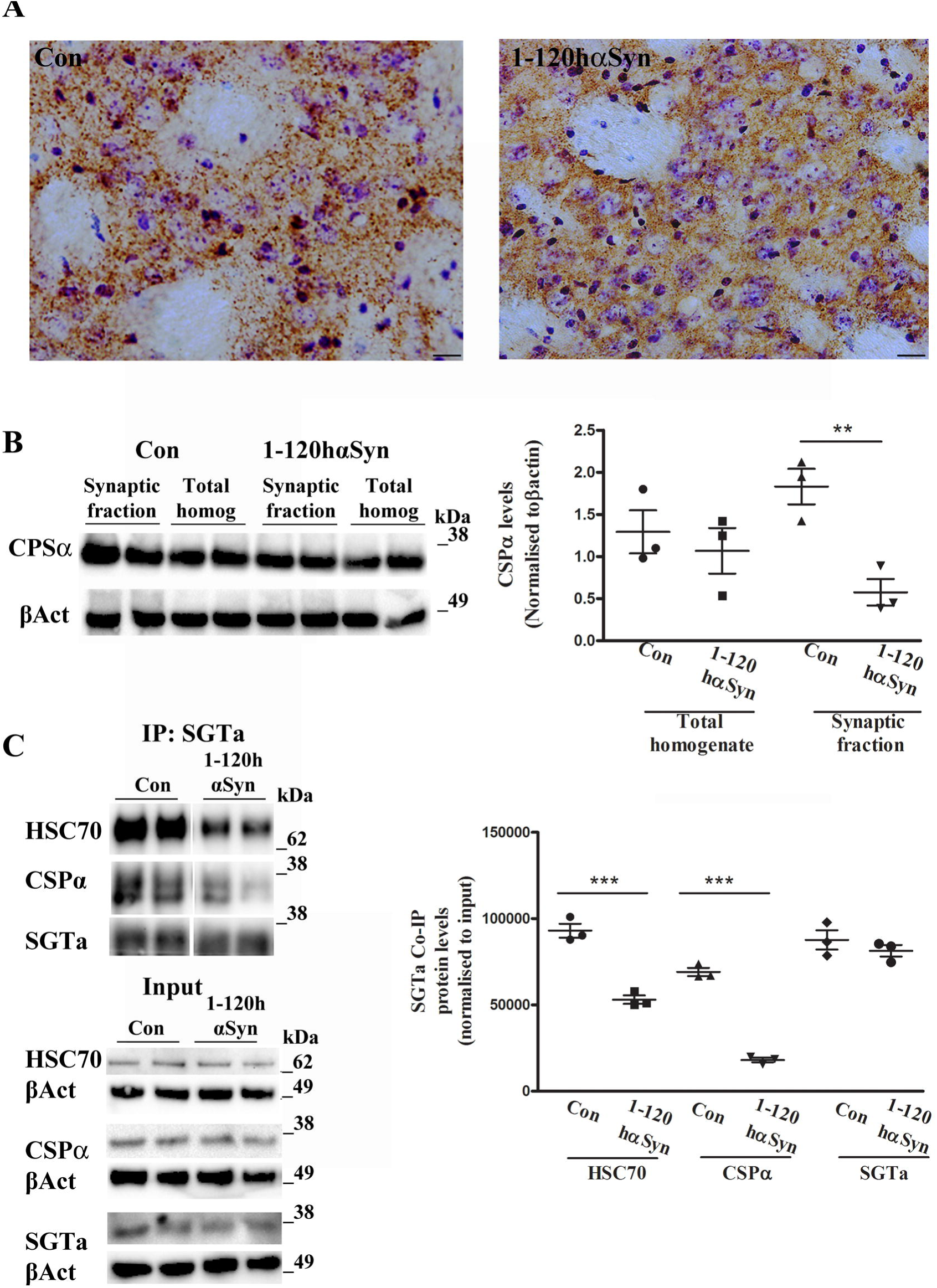
CSPα expression is altered in the striatum of 12 month-old 1-120hαSyn mice. **A** CSPα staining in striatal sections from controls and 1-120hαSyn mice. Note the loss of puncta intensity of CSPα in 1-120hαSyn mice compared to controls. Scale bar: 20 ¼m. **B** Left panel: Immunoblot for anti-CSPα and -βActin (βAct) in sequentially-extracted total homogenates and synaptic fractions from striata of control and 1-120hαSyn mice. Right panel: RI of CSPα levels normalised to βactin in total homogenates and synaptic fractions. Data are presented as mean±SEM of N=3 mice, **P<0.01 (Student’s t-Test). **C** Left panel; immunoblots of HSC70, CSPα, SGTa after co-immunoprecipitation with anti-SGTa antibody with non-hydrolysable ADP. CSPα and HSC70 complexes with SGTa are reduced in 1-120hαSyn mice compared to controls whereas input levels do not change (lower panel, CSPα, HSC70 and SGTa and correspondent βActin). Right panel; RI of HSC70, CSPα, SGTa relative to their input levels (total homogenates prior co-immunoprecipitation). Values in graph represent N=4-5 mice in 3 independent experiments ***P<0.001 (One-way ANOVA with Bonferroni’s multiple comparison test).

### CSPα rescues vesicle cycling impairment in PC12 cells expressing 1-120hαSyn

As previously reported expression of 1-120hαSyn in PC12 cells alters vesicle endo/exocytosis (Garcia-Reitbock *et al*., 2010). To determine whether CSPα could rescue the αsyn-related vesicle turnover impairment, PC12 cells stably expressing 1-120hαSyn were transduced with either an AAV6 vector encoding human CSPα (AAVCSPα), or an empty AAV6 control vector (AAVEV). In a control experiment, non–transfected PC12 cells were also treated with AAVCSPα or EV (Fig. 2).

**Figure 2.**
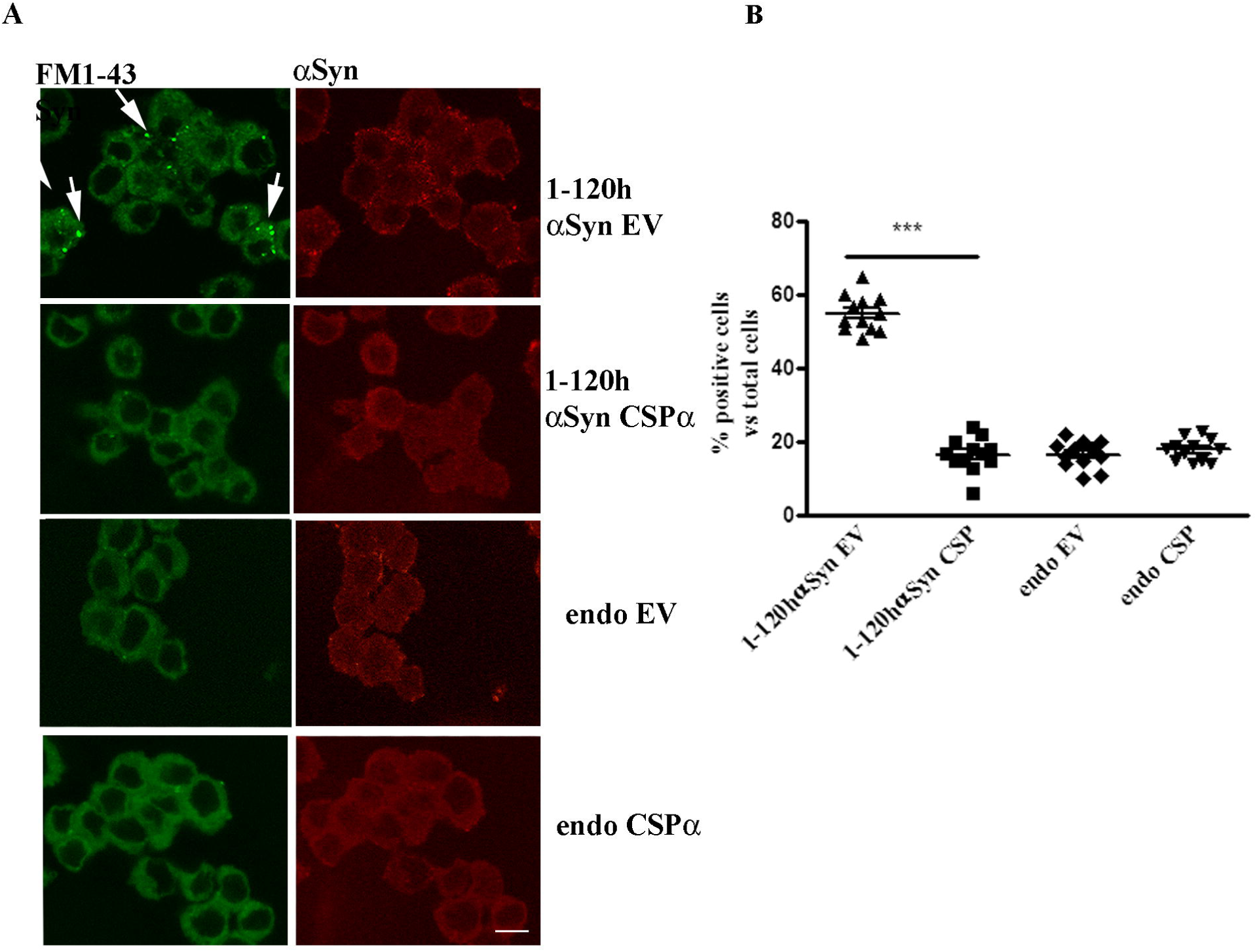
CSPα rescues the vesicle cycle impairment in PC12 cells expressing 1-120hαSyn. **A** FM1-43 dye fluorescence (left panels, green) and αsyn staining (right panels, red) in cells treated with empty vector (EV) or CSPα. Note the reduction in FM1-43 dye retention in cells stably expressing 1-120hαsyn treated with CSPα compared to their EV-treated counterpart (arrows). Treatment with CSPα or EV had no effect in non-transfected PC12 cells (endo). **B** Quantification of the number of cells that retained FM1-43 dye above basal levels in every treatment group. Values are mean±SEM of N=12 independent experiments ***P<0.001, Two-Way ANOVA Bonferroni’s multiple comparison test. Scale bar 10μm.

A cycle of vesicle uptake and release was induced by K^+^ in the presence of FM1-43 dye and the number of cells that internalised and retained the dye above a threshold value of fluorescence was counted. Fluorescence was increased in cells stably expressing 1-120hαSyn following K^+^ treatment, compared to un-transfected cells (endo) as previously shown (Garcia-Reitbock *et al*., 2010). Expression of CSPα in 1-120hαSyn expressing cells reduced the number of cells that retained FM1-43 while no effect was observed in cells treated with EV. Furthermore, CSPα or EV viral expression did not alter the release of FM1-43 from wild-type cells expressing endogenous αsyn (Fig. 2B) (cell number: 1-120hαsyn: EV =55±4.9, CSPα =16.6±4.6; endo: EV 16.9±3.6, CSPα 17.9±3). Thus, CSPα overexpression selectively rescues the synaptic vesicle cycle impairment caused by the expression of human truncated αsyn.

### CSPα delivery *in vivo* restores DA release

To investigate whether the effect of CSPα observed *in vitro* was present *in vivo*, we injected AAVCSPα and AAVEV into the SN of transgenic 1-120hαSyn mice which have a decrease in striatal DA release (Garcia-Reitbock *et al*, 2010) We first investigated the time-course of CSPα expression after injection in the SN of control mice with AAVCSPα and AAVEV and measured CSPα protein levels by immunoblotting in the striatum at 4, 6- and 8-weeks post-injection. A two-fold increase in CSPα expression in the striatum was present up to eight weeks post-injection (4 weeks: CSPα= 1.36±0.19, EV= 0.71± 0.13, 6 weeks: CSPα= 1.25± 0.08, EV= 0.46± 0.04; 8 weeks: CSPα= 1.22± 0.02, EV= 0.60± 0.08) (Supplementary Fig. 2). Therefore, AAVCSPα or AAVEV were injected in the SN of 1-120hαSyn and control 10 month-old mice and the striatal DA release was measured at eight weeks post-injection using *in vivo* microdialysis.

As previously shown, K^±^-evoked DA release was significantly reduced in untreated 1-120hαSyn mice compared to controls (Garcia-Reitbock *et al*., 2010) (fractions: 60, 80, 100 min normalised to peak control mice value (60 min); 0.47±0.045 vs 1±0.03 (60 min), 0.21±0.03 vs 0.63±0.18 (80 min), 0.12±0.03 vs 0.42±0.18 (100 min). Notably, expression of CSPα restored DA release levels in 1-120hαSyn mice back to the control values (0.98±0.06 fraction 60 min, 0.77±0.08 fraction 80 min) whereas the EV vector was ineffective (0.46±0.05 fraction 60 min, 0.29±0.05 fraction 80 min) (Fig. 3).

**Figure 3.**
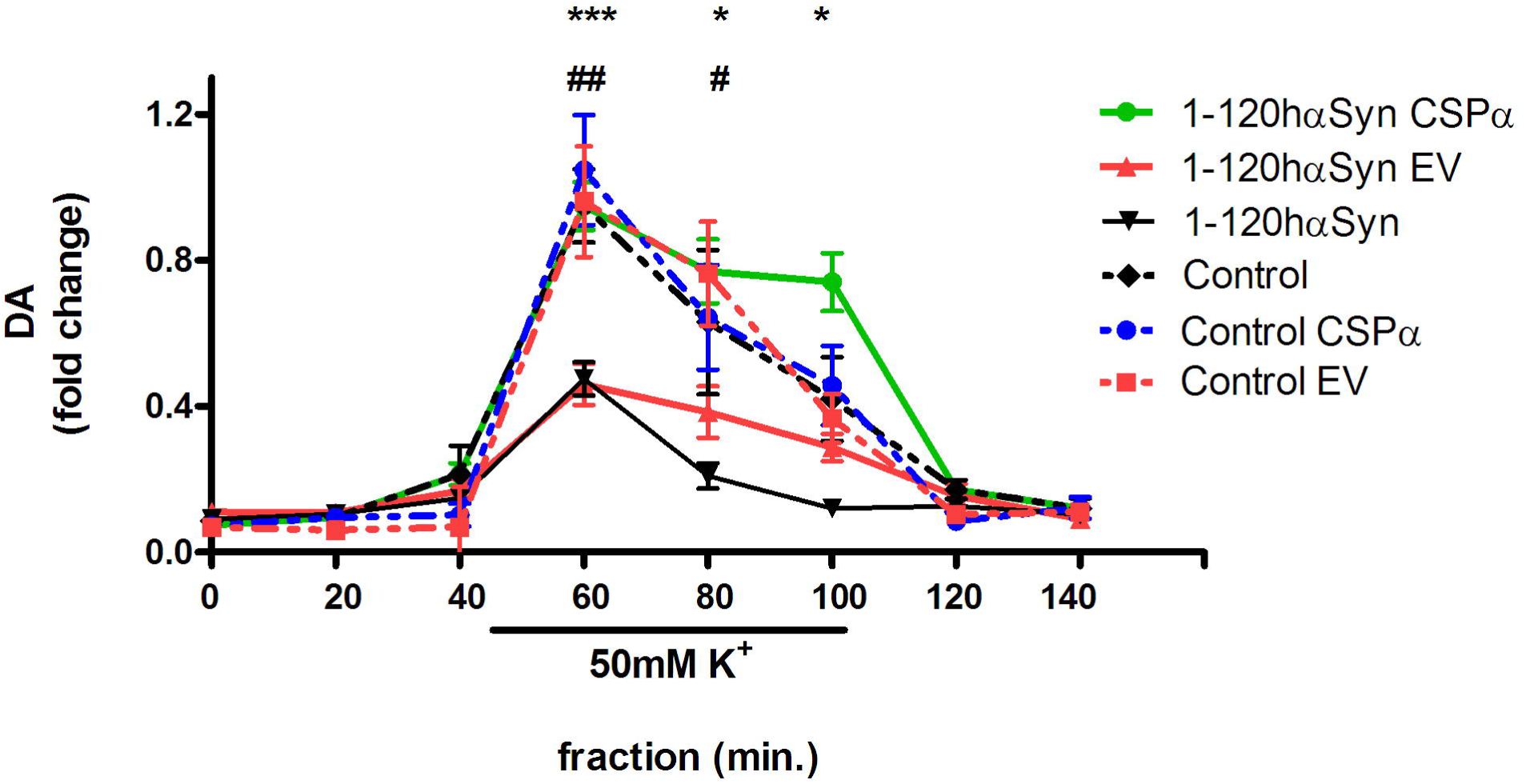
CSPα restore DA release impairment in 1-120hαsyn mice. Striatal DA release after infusion of 50mM KCl for 60 min during *in vivo* microdialysis. Twelve month-old 1-120hαSyn mice showed a significant reduction in DA release following KCl stimulation compared with controls (*** P<0.001, *P<0.05). Values show fold change compared to baseline fractions. Increase of CSPα following AAVCSPα injection restored DA release to control levels in 1-120hαSyn mice whereas treatment with AAVEV was ineffective (fractions: 60 min ## P<0.01, 80 min # P<0.05). No significant difference was observed in the 60- and 80-min fractions between CSPα and EV-treated or untreated control mice. Values are mean±SEM of N=5-7 1-120hαSyn and N=5-6 control mice per treatment group, two-way ANOVA, Bonferroni’s multiple comparison test.

The effect of CSPα was dependent on αsyn aggregation, because no difference was observed in control mice after treatment with CSPα or EV. (CSPα: 0.96±0.15 vs EV: 0.98±0.15 fraction 60 min, CSPα: 0.67±0.14 vs EV: 0.69±0.1 fraction 80 min) (Fig. 3).

These results show that increased expression of CSPα *in vivo* specifically rescues impaired DA release associated with 1-120 αsyn aggregation.

### CSPα reduces αsyn aggregates and increases αsyn monomers in the striatum of 1-120 hαSyn mice

To investigate whether CSPα increase had an effect on aggregation in 1-120hαSyn mice, immunostaining for αsyn was performed in striata of 12 months-old mice after microdialysis. Striata from EV-treated mice showed intensely stained αsyn aggregates in the neuropil, while the staining was reduced in CSPα-treated mice (Fig. 4A). To further characterise the αsyn species in CSPα animals, we used *d*STORM and compared the number of aggregated versus monomeric αsyn species at the nanoscopic level in both CSPα and EV-injected mice. In CSPα-injected mice the number of αsyn aggregates was reduced compared to EV-treated mice (aggregate number relative units (RU)=% of all species that are aggregated: CSPα= 66± 2%, EV=78±2%) (Fig 4B).

**Figure 4.**
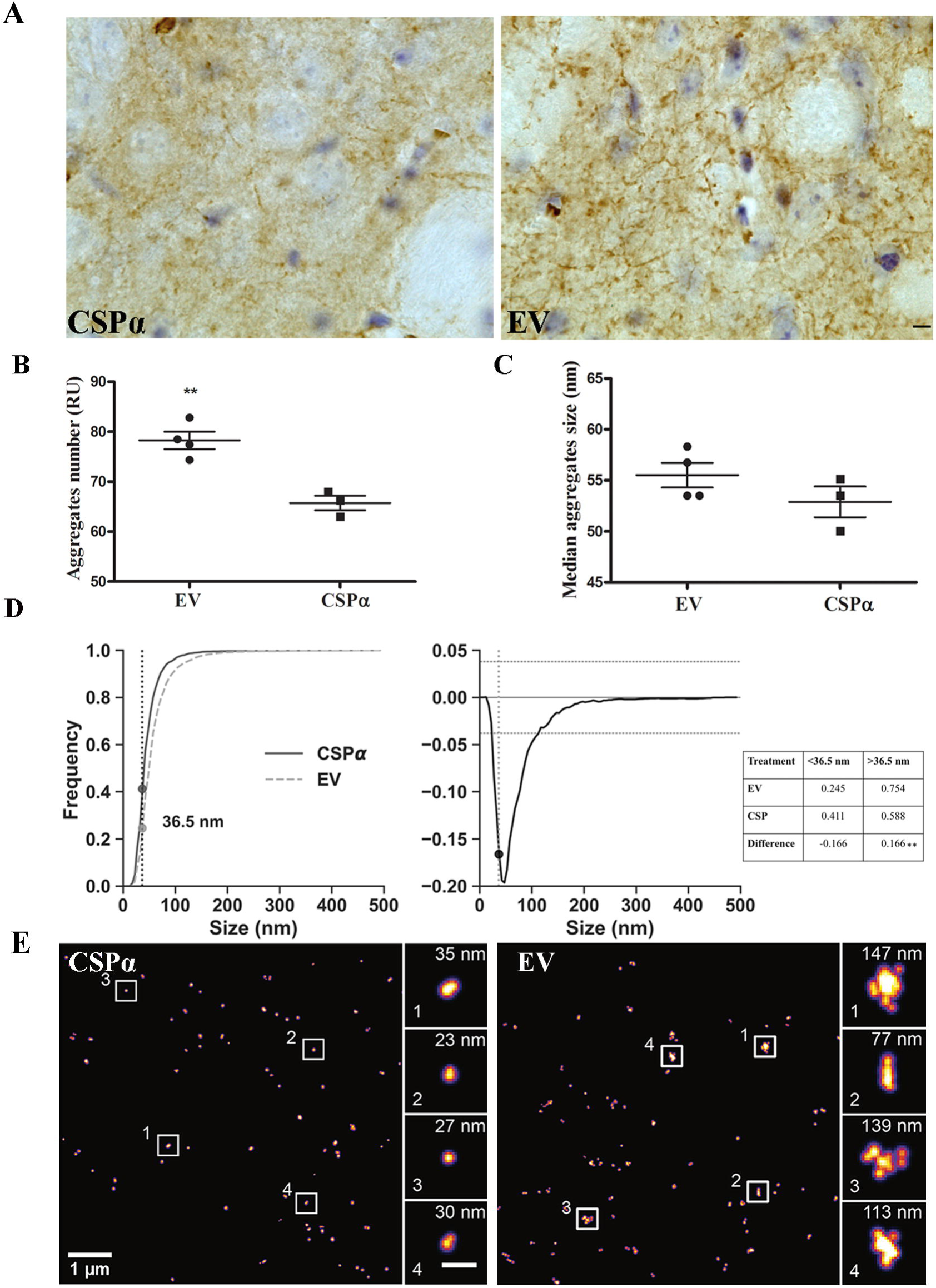
CSPα reduces αsyn aggregates and increases αsyn monomers in the striatum of 1-120hαSyn mice. **A** αSyn immunostaining in the striata of CSPα and EV treated 1-120hαsyn mice. Scale bar 10µm. **B** Aggregate number (RU) versus monomeric αsyn in AAVEV (EV) and AAVCSPα (CSPα)-treated mice based on *d*STORM analysis using anti-αsyn Syn1 antibody. A significant reduction in the number of aggregates versus monomeric αsyn is present after AAVCSPα injection. Two-tailed Student’s t test **p≤0.01; N=4 mice per group. **C** Median size of αsyn aggregates showing no significant difference between EV-and CSPα-injected mice. Two-tailed Student’s t test p≥0.01, N=4 mice. **D** Left panel, Cumulative histograms for αsyn species distribution in EV- and CSPα-treated mice and difference between the two distributions. Right panel, Frequency values table of αsyn species at the 36.5 nm intercept in EV- and CSPα-treated mice. Kolmogorov-Smirnov test **P<0.01, number of measured species EV: 4583, CSPα: 2574. **E** Representative *d*STORM images of αsyn staining in CSPα or EV injected 1-120hαsyn mouse striatum. Note the difference in size of αsyn aggregates, (single αsyn aggregates enlarged in boxed areas scale bar insets =200 nm).

We then analysed the difference in aggregate size distribution between EV- and CSPα-injected mice. No difference in the median size of αsyn aggregates between the CSPα-injected and EV injected group was found, however CSPα-injected mice exhibited a significant 16.6% reduction in αsyn aggregates (>36.5 nm) and conversely, an increase in monomeric species (<36.5 nm) (Fig 4C, D and Table) indicating that CSPα reduced the number of αsyn aggregates and increased the number of αsyn monomeric species.

## Discussion

αSyn is predominantly localised at the synapse where it is involved in SNARE complex assembly and synaptic vesicle turnover (Iwai *et al*., 1995; Nemani *et al*., 2010; Burre *et al*., 2010, 2014; Diao *et al*., 2013; Vargas *et al*., 2017; Longhena *et al*. 2019). CSPα is also at the synapse where it assists folding of client proteins involved in neurotransmitter release, exo/endocytosis and SNARE complex formation (Tobaben *et al*., 2001; Chandra *et al*., 2005; Zhang *et al*., 2012). αSyn is functionally interconnected with CSPα as shown by its ability to rescue the neurodegeneration characteristic of CSPα-KO mice (Chandra *et al*., 2005) but whether CSPα is affected by αsyn aggregation is not clear. Here we show that CSPα immunoreactivity is reduced in the striatum of 12 month-old 1-120hαSyn mice compared to controls. This decrease was specifically in the synaptosomal fraction indicating that CSPα reduction is localised where αsyn aggregates. As a synaptic co-chaperone, CSPα binds to HSC70 and the adapter protein SGTa to regulate vesicle fusion. By co-immunoprecipitation we found that CSPα/HSC70 complexes with SGTa are reduced in the striatum, effect that may be attributed to CSPα reduction. These findings indicate that αsyn synaptic aggregation affects CSPα levels and function. In order to determine whether an increase of CSPα could be beneficial and restore vesicle cycle and DA release, we tested this first in PC12 cells expressing 1-120hαsyn and found that indeed their abnormal vesicle cycle was restored. This prompted us to investigate whether an increase of CSPα *in vivo* in the striatum of 1-120hαSyn mice could restore DA release. Using microdialysis we found that CSPα expression restored the striatal DA release reduced by synaptic αsyn aggregation, and *d*STORM super-resolution microscopy performed in tissue after microdialysis, indicated a reduction in the number of αsyn aggregates and a concomitant increase in monomeric species. It is unclear whether CSPα prevents monomeric αsyn aggregation or contributes to dissociation of the αsyn aggregates. Both HSC70 small chaperones and other DNAJs have been shown to affect αsyn aggregation (Pemberton *et al*., 2011; Whiten *et al*., 2018,*a*), in our model, CSPα could contribute to facilitate HSC70 ATPase activity as the burden of misfolded αsyn increases. Although it cannot be excluded that the increase in CSPα per se could improve dopamine release, the fact that no change was observed in control mice would support that CSPα acts by affecting αsyn-related alterations. The increase in monomeric αsyn is similar to what we observed in MI2 transgenic mice after anle138b treatment (Wegrzynowicz *et al*., 2019), whether CSPα beneficial effect is due the reduction of aggregates or the increase of monomeric αsyn remains to be determined.

In conclusion, our data reveal for the first time that αsyn aggregation impairs expression of synaptic CSPα and formation of its functional complexes with HSC70 and that increase of CSPα reduces synaptic αsyn aggregates and increases monomeric αsyn rescuing DA release impairment. These results point to CSPα and related chaperones as therapeutic avenues for restoring normal synaptic function in early PD.

## Acknowledgements

We wish to thank Prof RD Burgoyne for the CSPα cDNA, Dr E. Dimou and Dr J. McColl for advice on *d*STORM imaging and Dr MP Fransen and Dr G Vivacqua for critically reading the manuscript.

## Abbreviations

CSPα: cysteine string protein
KO: knock-out
1-120hαSyn: 1-120 truncated human αsynuclein
PD: Parkinson’s disease
SNARE: soluble N-ethylmaleimide sensitive fusion attachment protein receptor

## Supplementary Figure Legends

**Supplementary Figure 1 HSC70, SGTa and VAMP2 expression in the striatum A** (Left panel) Representative blots of synaptosomal fraction of HSC70, correspondent βActin and normalised RI values (right panel, Control (Con) 0.29±0.05, 1-120hαSyn 0.42± 0.1). **B** (Left panel) Representative blots of synaptosomal fraction of SGTa, correspondent βActin and normalised RI values (Right panel, Con 1.32±0.06, 1-120hαSyn 1.28± 0.08). **C** (Left panel) Representative blots of VAMP2 and correspondent βActin in control and 1-120hαSyn mice from synaptic and total brain homogenates fractions. (Right panel) VAMP2 levels quantification (Synaptic fraction RI; Con 0.72±0.07, 1-120hαSyn, 0.65±0.03; total homogenate; Con 0.25±0.06, 1-120hαSyn 0.34± 0.03) data are mean±SEM of N=3 mice per group, two-tailed Students t test for data in A an B, one-way ANOVA for data in C.

**Supplementary Figure 2 Time course of CSP**α **expression following injection of AAVCSP**α **and AAVEV in mouse substantia nigra. A** Immunoblots of CSPα and βActin in striatal extracts at 4, 6, 8 weeks post AAV injections in the SN. **B** Densitometry of CSPα levels at 4, 6 and 8 weeks post AAV injection. Data are mean±SEM of N=3 mice per group, Two-way ANOVA with Bonferroni’s multiple comparisons test, 4wks ** P<0.01, 6 wks ***P<0.001, 8 wks ** P<0.01.

