## Supplementary figures and images for "CSPα reduces aggregates and rescues striatal dopamine release in αsynuclein transgenic mice"

### Supplementary Fig 1

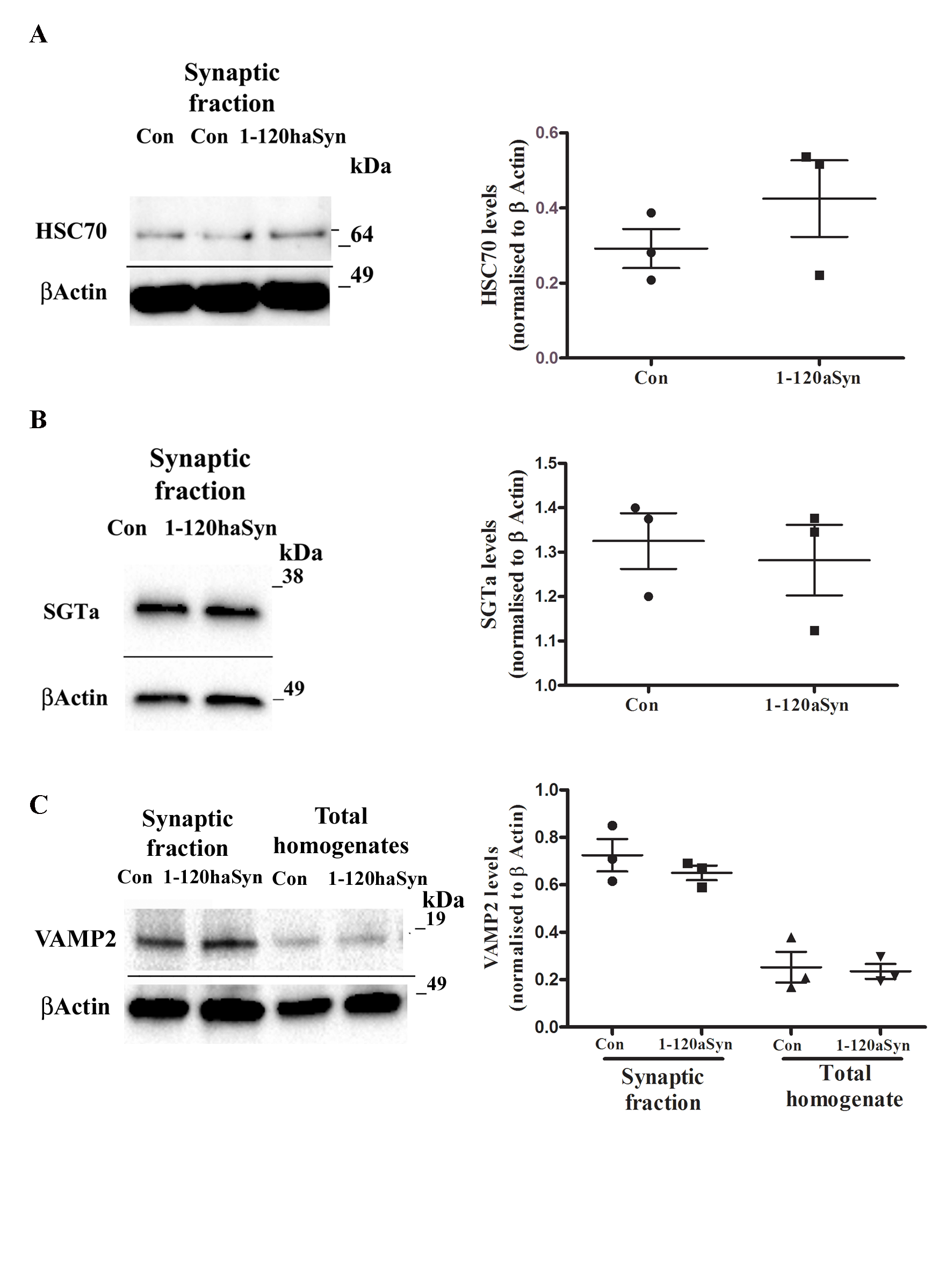

### Supplementary Fig 2

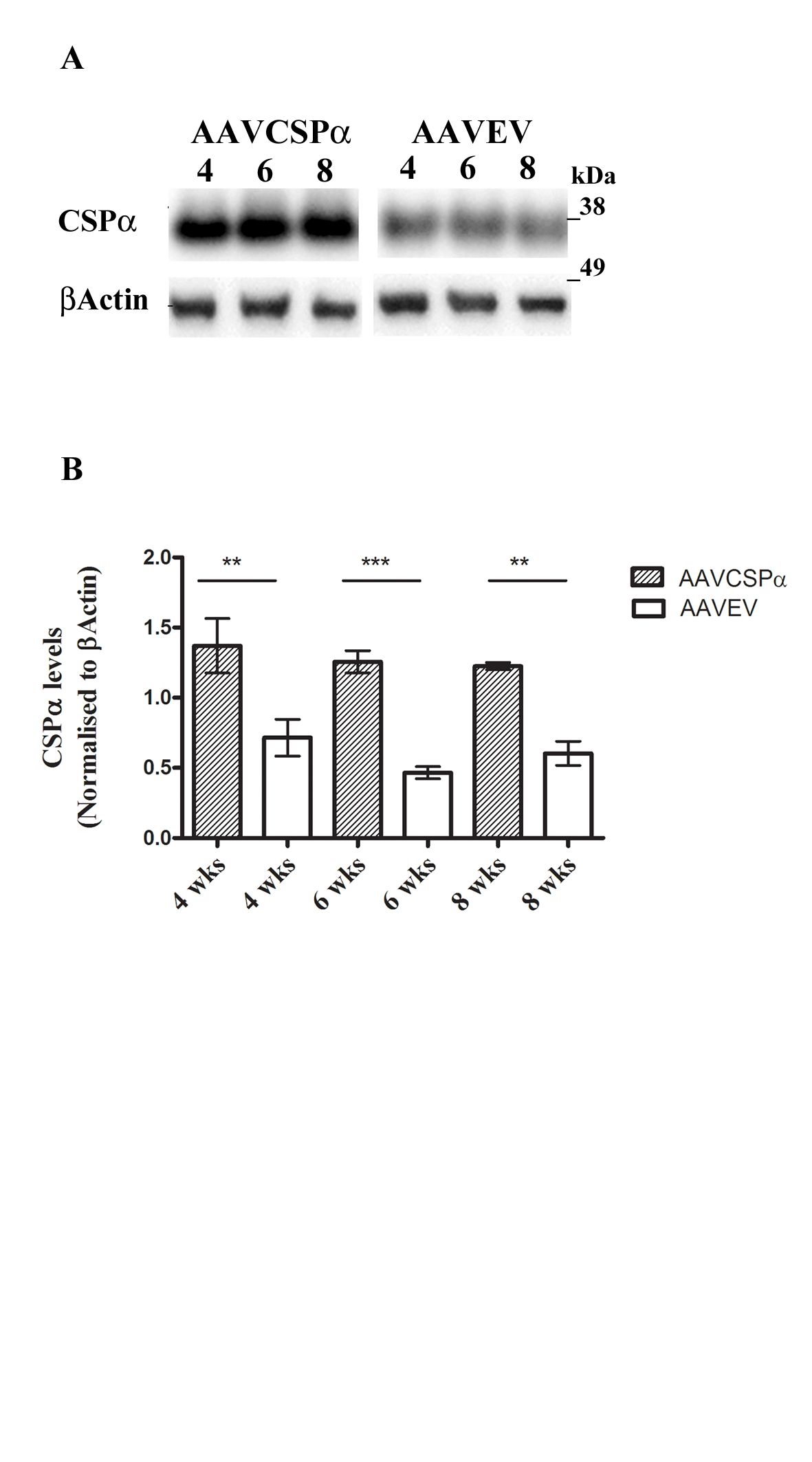
